# The black goby *Gobius niger* Linnaeus, 1758 in the Marchica lagoon (Alboran Sea, Morocco): Spatial-temporal distribution and its environmental drivers, and the site-related footprint

**DOI:** 10.1101/2022.10.02.510494

**Authors:** Amal Lamkhalkhal, Mohamed Selfati, Imane Rahmouni, Nassir Kaddouri, Bouabid Badaoui, Antoine Pariselle, Abdelaziz Benhoussa, Marcelo Kovačić, Nikol Kmentová, Maarten P.M. Vanhove, Hocein Bazairi

## Abstract

Fishes belonging to Gobiidae are well represented in the Marchica lagoon on the Moroccan Mediterranean coast, both in terms of species richness and abundance, with the black goby, *Gobius niger* Linnaeus, 1758, being the dominant species. The present study aims to examine (1) the spatial and temporal distribution of *Gobius niger* and its environmental drivers in the lagoon and (2) the potential lagoon-related footprint using morphometric, genetic and parasitological proxies.

Systematic monthly sampling covering the whole lagoon basin performed between October 2015 and September 2016 revealed year-long presence of *G. niger* throughout the lagoon with significantly low densities in winter. The higher abundances were recorded in the shallow bottoms of the lagoon inner margins on a variety of substrates (mud, muddy-sand, sandy-mud and fine sand) mostly covered by macroalgae and/or seagrass meadows. Depth, that has to be seen as a variable that acts in concert with other factors such as temperature, vegetation cover and sediment as the Mean Grain Size, seems to be the most important predictor, explaining the distribution of *G. niger* in the lagoon, with a trend of increasing abundance towards shallower stations.

Comparison of black goby populations from the Marchica lagoon with their counterparts from the adjacent Mediterranean coast of Morocco revealed that specimens caught at sea are of a bigger size compared to the lagoon population. Of the 180 gobies investigated, not a single one hosted the parasites we targeted in the parasitological approach, monogenean flatworms. The absence of population structuring, low genetic diversity and presence of common haplotypes indicate no apparent restriction in the gene flow between the two populations. Therefore, the observed morphometric differences seem to be due to the external environmental conditions rather than genetic differences.

*Gobius niger* plays a key eco-trophic role by providing a link between benthic invertebrates and large predators. The shallow beds of the lagoon, where the species is abundant, are key habitats in the Marchica lagoon and need to be considered in all management plans aiming at the conservation of biodiversity and ecological processes.

## Introduction

Coastal lagoons are aquatic ecosystems at the terrestrial and marine interface, occupying approximately 13% of the world’s coastline (Kjerfve, 1994). Due to the multiple ecosystem services they provide (Levin *et al*., 2001) (*e.g*. shoreline protection, fisheries resources, nursery area, etc.), lagoons are considered as one of the most valuable coastal habitats on the planet (Pérez-Ruzafa *et al*., 2019). However, combined natural and man-made stressors make them among the most heavily exploited and threatened natural systems worldwide (Eisenreich, 2005; Newton *et al*., 2018).

Fish play a fundamental role in the ecological processes, through trophic relationships with other biotic components (Stein *et al*., 1995; Vanni, 2002), and are essential for the functioning and resilience of lagoon ecosystems (Koutrakis *et al*., 2005; Franco *et al*., 2006; Aliaume *et al*., 2007. They are highly valuable for the local human population as food supply and for job creation (Holmlund & Hammer, 1999; Lopes & Videira, 2013; Newton *et al*., 2014). Moreover, fish are relevant biotic indicators to survey biodiversity and ecological status of the ecosystem (Whitfield & Elliott, 2002; Breine *et al*., 2010). Therefore, reliable scientific data on the fish fauna and abiotic components of lagoon ecosystems are of particular importance for effective management (Vasconcelos & Galyean, 2007), ensuring the sustainability of ecosystem functions and services.

With 2,949 currently recognized species, Gobiidae is the most species-rich family of fishes, (Fricke *et al*., 2023; Renoult *et al*., 2022). Gobies are found both in marine and freshwater environments (Renoult *et al*., 2022). They show highest species richness in warm temperate and tropical seas, on the continental shelf, mostly in the shallow part. Generally, they are small and short-lived, and most of them live discreetly on the substrate or hidden in various types of hidden spaces. Among them are epibenthic, hyperbenthic and cryptobenthic species, while some gobiid species are nektonic (Kovačić & Patzner, 2011). Gobies have a crucial trophic function by linking benthic invertebrates to larger predatory fish (Casabianca & Kiener, 1969; Miller, 1979; Raffaelli *et al*., 1989). Despite their low commercial value, gobies play an importantrole as food resources for many commercially important species.

In the Mediterranean basin, 78 species of Gobiidae are known to occur currently (Kovačić *et al*., 2022; Mavruk *et al*., 2022). In the Mediterranean, 249 fish species were listed as inhabiting estuaries and/or lagoons including both resident (euryhaline species, that live out their entire lifecycle inside lagoons and estuaries) and migratory (species that, after spending time in lagoons, are obliged to return to their native marine or river environment to complete their life cycle) fishes (Kara & Quignard, 2019). Besides, Gobiidae (at least 11 species) constitutes, with Syngnathidae (at least ten species), the most represented family of resident fish in Mediterranean lagoons (Kara & Quignard, 2019).

The Marchica lagoon (35.156944° / -2.845278°), situated on the Moroccan Mediterranean coast, is known to host a diverse fish fauna supporting important fishing activities (Selfati, 2020). Since the first inventory in 1911 (Oden, 1914), Gobiidae has been found to be well represented in the Marchica lagoon, both qualitatively and quantitatively, with the black goby, *Gobius niger* Linnaeus, 1758, as the dominant species in the fish fauna (Selfati *et al*., 2020). However, very little information exists on the ecological characteristics of this species, in particular its spatio-temporal structure in relation to environmental factors.

Resident animals, such as some gobies, are the most suitable to be used as an indicator of lagoon environmental conditions (Bortone *et al*., 2005). Their presence and abundance may provide important indications on the conservation status of coastal lagoon habitats (Facca *et al*., 2020). Considering that the black goby *G. niger* is the most common species of gobies in the Marchica lagoon, that it is a resident fish in coastal lagoons (Franco *et al*., 2008a, 2008b, 2012; Kara & Quignard, 2019; Selfati *et al*., 2019) and that its lagoon population has its counterparts of the same species along the adjacent Mediterranean coast of Morocco, our study aims to (1) analyse the spatial and temporal distribution of *G. niger* and its environmental drivers using a systematic sampling covering the whole lagoon basin, and (2) examine the potential lagoon-related footprint *i.e*. whether the supposed sedentary population of the black goby in the Marchica lagoon differs from black gobies from the adjacent Mediterranean coast of Morocco considering morphometric, genetic and parasitological proxies.

## Materials and Methods

### Study area

The Marchica lagoon (Fig. 1) (35.156944° / -2.845278°), also called the lagoon of Nador, is one of the largest coastal lagoons in the Mediterranean (115 km², 25 km long and 7.5 km wide) and the only one on the Mediterranean coast of Morocco (Selfati, 2020). The maximum depth is approximately 8m and the lagoon is separated from the Mediterranean Sea by a 25-km-long sandbar (Lido), with one artificial opening (300 m wide and 6 m deep) that allows water exchange. Despite its ecological (Site of Biological and Ecological Interest since 1996; RAMSAR site since 2005) and socio-economic (mainly artisanal fisheries) values, the lagoon is under pressure from a complex mixture of human-mediated stressors (urbanization, pollution, overfishing, tourism, and wastewater, among others) (Selfati *et al*., 2017; El Kamcha *et al*., 2020).

**Fig. 1:**
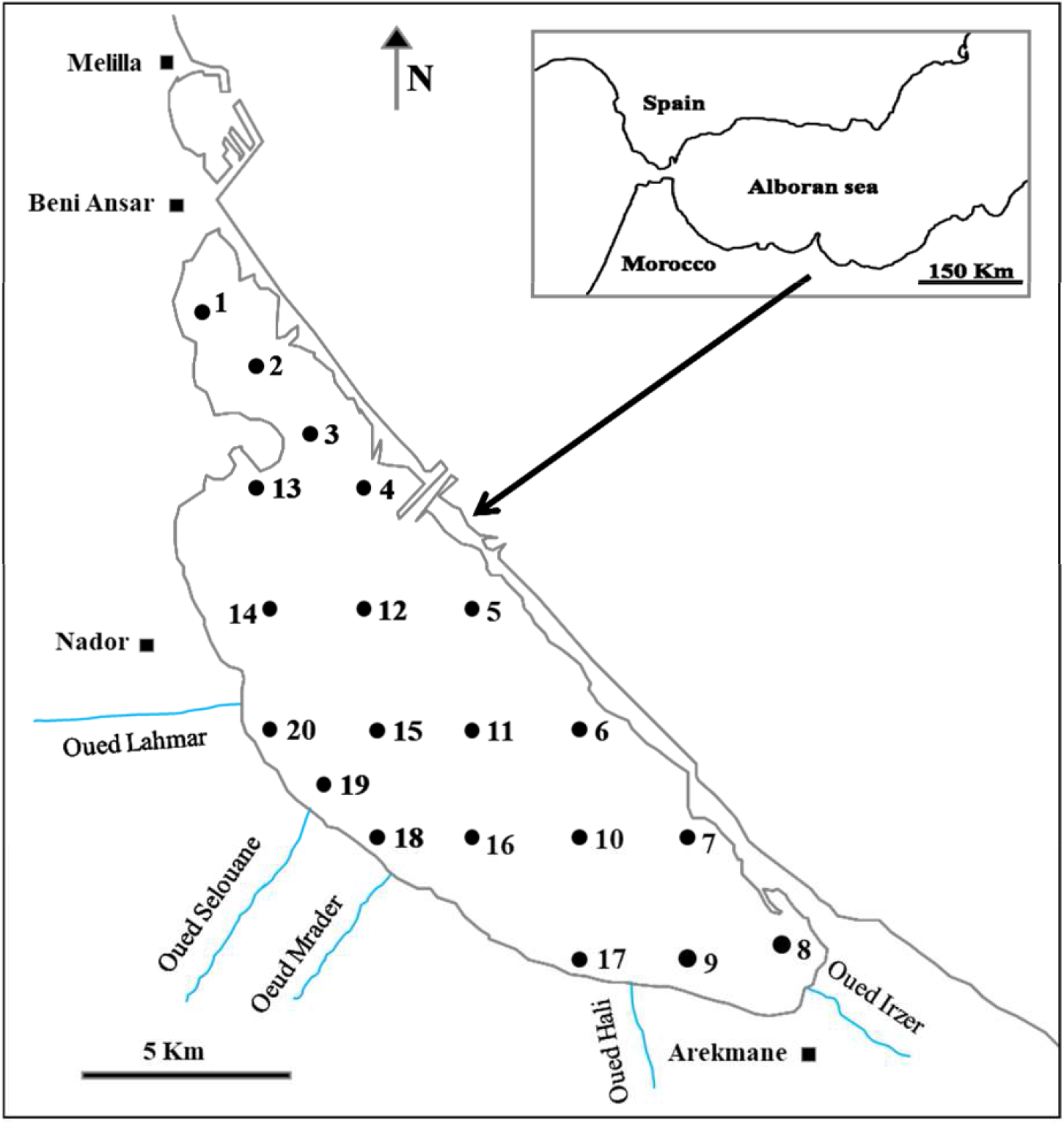
Map showing the geographical localization of the Marchica lagoon and the sampling stations of *Gobius niger*.

### Sampling design and environmental data

To examine the spatial and temporal variations in species abundance of *G. niger* and its environmental drivers in the Marchica lagoon, data on monthly abundances was extracted from a scientific monitoring of the fish fauna in the lagoon between October 2015 and September 2016. The monitoring was carried out according to an optimized network of 20 stations (S1-S20) covering the whole lagoon (Fig. 1). The collecting gear was a purse seine of about 110 m in length and 11 m in height, with a mesh size of 6 mm. The catches were expressed by units of effort corresponding to the surface sampled: the deployment of the seine describes a circle on the surface of the water with an estimated radius of 16 meters allowing to deduce the encircled surface equivalent to 800 m². Environmental parameters were measured at each station. Water temperature (T), salinity (S) and conductivity (Cond, mS/cm) were measured *in situ* using a conductivity meter ‘‘Cond 315i/SET’’, and depth (D) was recorded using an LCD Digital Sounder (HONDEX PS-7). The water pH was measured in the laboratory from water samples collected in the field using a pH meter ‘‘IONOMETER-EUTECHINSTRUMENTS-CYBERSCAN-PH-510’’. The assay of suspended matter (SM, mg/l) in water samples was performed in the laboratory by the 0.45-μm membrane filtration method. The percentage of vegetation cover (VC: combined seagrass and macroalgae) and the nature of the substratum expressed as mean grain size (MGS, *u*m) were also derived from the literature (Najih et al., 2016, Najih et al., 2017). Data were obtained by projection of the fish sampling points on the vegetation cover map and on the map of distribution of surface sediment facies.

To examine the potential lagoon-related footprint, *i.e*. whether the supposedly sedentary population of the black goby in the Marchica lagoon differs from black gobies from the adjacent Mediterranean coast of Morocco considering morphometric, genetic and parasitological proxies, 120 black gobies from the Marchica lagoon and 60 ones from the adjacent Mediterranean coast of Morocco were all collected as bycatch by fishing boats in July 2020. All the specimens were stored in separated plastic bags and were transported in a portable freezer (Engel MT45), then stored in the laboratory in a freezer (−20°C) for further study.

After thawing, the fish were numbered, labelled and photographed. A piece of the pectoral fin of each fish was taken and kept in an Eppendorf tube filled with 96% ethanol for molecular characterization.

### Species identification

The first samples of the species were identified by minimum combination of characters that positively identify the specimens of *G. niger* among species of Gobiidae in the Mediterranean (Kovačić, 2020): 1) Suborbital sensory papillae of the head lateral line system without suborbital row *a*; 2) All three head canals of the head lateral line system present; 3) Anterior dorsal row *g* of sensory papillae ends behind or on lateral end of row *o*; 4) Six suborbital transverse rows *c* of sensory papillae; 5) Anterior oculoscapular head canal with pore *α* at rear of orbit; 6) Oculoscapular row *x¹* of sensory papillae ending forward behind pore *β*; 7) Longitudinal scale count <50; 8) Predorsal area scaled: 9) Suborbital row *d* of sensory papillae continuous.

Identification of every later sample was based on a simpler determination inspired by those provided by Brownell & Collignon (1978), Bauchot & Pras (1980) and Bauchot (1987).

### Spatial and temporal distribution of Gobius niger in the Marchica lagoon and environmental drivers

Spatial and temporal variations in abundance of *G. niger* (expressed as densities per 800 m²) were illustrated on maps using 11 classes of abundance based on the Sturges rule (Sturges, 1926). The spatial pattern of abundances (expressed as abundance per month and per station) of *G. niger* in the Marchica lagoon was explored, to identify affinity groups of stations, using a hierarchical cluster analysis, conducted on a transformed (four root) abundances similarity matrix based on the Euclidean distances. Then, the Analysis of similarities (ANOSIM) non-parametric test was performed to assess the level of significance of the groups of stations identified. Differences between sampling stations and between seasons (winter: December, January and February; spring: March, April and May; summer: June, July and August; autumn: September, October and November) were tested with a two-way crossed PERMANOVA design.

Distance-based linear modelling (DISTLM) was performed to identify the key environmental drivers of the black goby’s distribution pattern in the Marchica lagoon. The best overall model was selected using the BEST selection procedure with the Akaike Information Criterion (AIC) in order to reveal the significant variables influencing the observed patterns in spatial abundance (Akaike, 1973; Anderson *et al*., 2008). A distance-based redundancy analysis (dbRDA, Legendre & Anderson, 1999; McArdle & Anderson, 2001) was used to illustrate graphically the results. The draftsman plots, and the associated correlation matrix between all pairs of variables, were examined for evidence of collinearity (Clarke *et al*., 2014).

All the multivariate analyses were performed using the PRIMER 6 software (Clarke & Gorley, 2005).

### Characterisation and comparison of goby populations from the Marchica lagoon and adjacent Mediterranean Sea

#### Morphometric analysis

Based on literature (Gaamour *et al*., 2001), six morphometric characters were measured on each specimen with a 1 mm resolution using a Vernier calliper: total length (TL), standard length (SL), head length (HL), snout length (SnL), body height (BH) and eye diameter (ED). Moreover, five meristic characters were considered: number of rays in the first dorsal fin (DF1), number of rays in the second dorsal fin (DF2), number of rays in the anal fin (AF), number of rays in the pectoral fin (PF) and number of rays in the ventral fin (VF) (Gaamour *et al*., 2001) (Fig. 2). Statistical analyses (Student’s t-test) were performed to detect differences in term size between specimens from two locations (Marchica lagoon and adjacent Sea).

**Fig. 2:**
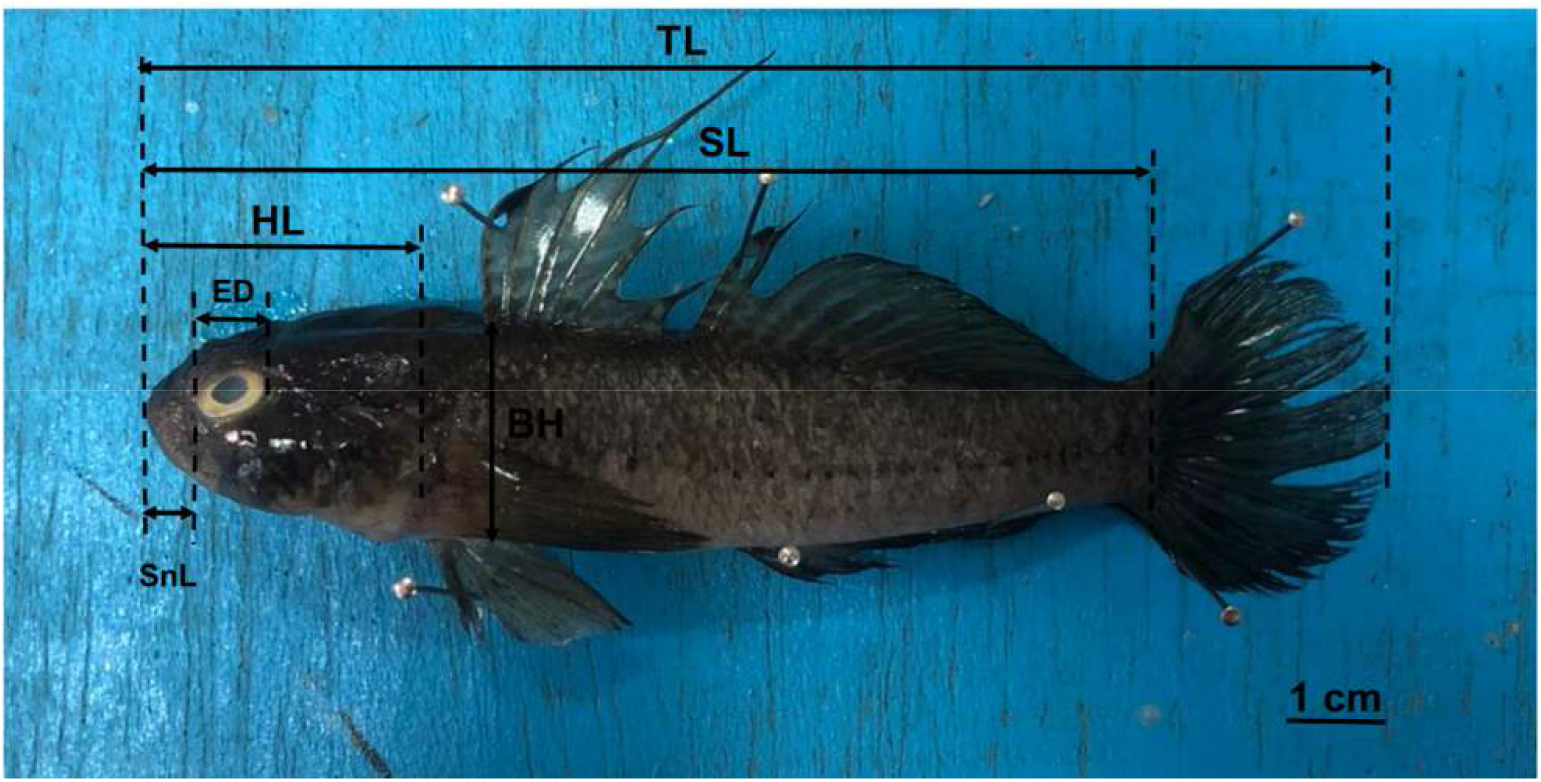
Picture of *Gobius niger* from the Marchica lagoon showing the main measurements taken: total length (TL), standard length (SL), head length (LT), snout length (SnL), Body height (BH) and eye diameter (ED).

In order to see if there is a stratification based on origin of *G. niger*, a Principal Component Analysis (PCA) was performed using morphometric and meristic characters, after a transformation (Log_10_X+1) of the raw measurements in order to linearize the allometries (Huxley, 1932) and to roughly equalize the variances (Jolicoeur, 1963).

All morphometric and meristic analyses were performed in the PAST v4.03 software (Hammer, Harper & Ryan, 2001).

### Genetic differentiation

DNA extraction of 120 specimens (60 from the Marchica lagoon and 60 from the adjacent Mediterranean coast) was performed according to the protocol of Aljanabi & Martinez (1997): approximately 50μg of pectoral fin fragment was digested at 55 °C overnight with 20μl of proteinase *K* (20 mg/ml) and 180μl of extraction buffer (0.4M NaCl, 1M Tris, 2 mM EDTA and 40 µl of 20 % SDS). The extracted DNA was suspended in 150μl of sterile double-distilled water and stored at -20°C until amplification by PCR. We targeted a fragment of the 16S rRNA gene, since mitochondrial ribosomal sequences are well-represented in the genetic literature on Mediterranean and European gobies. As they capture interspecific and intraspecific diversity in these fishes, they can be considered potential barcoding markers (Vanhove *et al*., 2012; 2016; 2022 and references therein). Amplification of the 16S rRNA gene was performed in a final volume of 20μl containing: 4μl buffer (10× Standard Taq Reaction Buffer), 1μl of 10 mM dNTPs, 0.8μl of 10μM forward primer 16SH (5’-CGCCTGTTTATCAAAAACAT-3’), 0.8μl of 10μM reverse primer 16SL 5’-CGCCTGTTTATCAAAAACAT-3’) (Palumbi *et al*., 1991), 0.4µl (2 units) of *Taq* polymerase, 1μl of genomic DNA and 12 of nuclease free water. Amplification reactions were performed in a thermal gradient PCR according to the following program: initial denaturation at 94°C for 3 min followed by 40 cycles each with denaturation for 30 sec at 94°C; a hybridization for 30 sec at 55°C and an elongation phase for 1 min at 72°C and at the end a final elongation for 10 min at 72°C. The PCR products were checked on 1% agarose gel and sent to the National Center for Scientific and Technical Research (CNRST) in Rabat; then they were sequenced by a Genomix sequencer (MGX) using the same forward and reverse primers as for the PCR.

Each DNA sequence obtained in both directions was cleaned and checked in MEGA X (Kumar *et al*., 2018) to assemble the corresponding consensus sequence. The obtained sequences were aligned with the CLUSTAL W algorithm (Thompson *et al*., 1994) and then each sequence was blasted (Altschul *et al*., 1990) with the sequences of *Gobius niger* 16S available in NCBI GenBank to check for possible matches.

To make sure that all of our 16S sequences, both the newly obtained Moroccan sequences, and those we include from other studies, cluster monophyletically, without representatives of other species, a phylogenetic tree was built including all other species of *Gobius* that are represented on GenBank by the targeted fragment of the 16S rRNA gene (see Supplementary material Table S1). The phylogenetic tree was constructed with MEGA X using the Neighbor Joining algorithm and the Kimura 2-P substitution model with 1000 bootstrap replicates.

All sequences were confirmed to belong to *G. niger* (Supplementary material Figure S1) and hence were included in a maximum parsimony analysis in the software package FITCHI (Matschiner, 2016) in order to visualize the genealogical relationships across the geographical range of *G. niger*.

DNASP v6.12.03 (Rozas *et al*., 2017) was used to calculate molecular diversity indices: number of segregating sites (K), number of haplotypes (H), haplotype diversity (h) and nucleotide diversity (π). Moreover, Fu, Li’s F and Tajima’s neutrality tests were performed to check for possible selection or change in population demography. Pairwise F_ST_ values were calculated in ARLEQUIN v3.5 (Excoffier & Lischer, 2010) using 100 permutations to express the degree of genetic differentiation among sets of individuals.

A matrix correlation analysis (Mantel test; Mantel, 1967) permuting a morphological distance matrix against a genetic distance matrix was performed using the R v3.5 software package ape (Paradis *et al*., 2004), running 1000 permutations.

### Parasitological screening

Each specimen was placed in plastic bags with tap water and then the bags were vigorously shaken to detach the parasites from the skin, then the external surface of skin, fins and the holding water were examined. Gill arches on the right side of each specimen were removed through ventral and dorsal section, placed in a petri dish, and rinsed with a rinsing bottle filled with tap water. Then water and gill arches were examined under a stereoscope (Wild M8).

## Results

### *Spatial and temporal distribution of* Gobius niger *in the Marchica lagoon and its environmental drivers*

Cluster analysis of *G. niger* abundances in the 20 sampled stations in the Marchica lagoon separated the samples in two significant groups (ANOSIM, P < 0.05), labelled as G1 and G2 (Fig. 3). The group G1, where abundances were high (mean= 95.44±120.44), concerns the NW (S1 and S2), the SE extremities of the lagoon (S7, S8 and S9) and the west, inner coast around Oued Selouane (S18, S19 and S20). The group G2 corresponds roughly to the center of the lagoon and the lagoon entrance where the goby was overall less abundant (mean= 9.91±18.01).

**Fig. 3:**
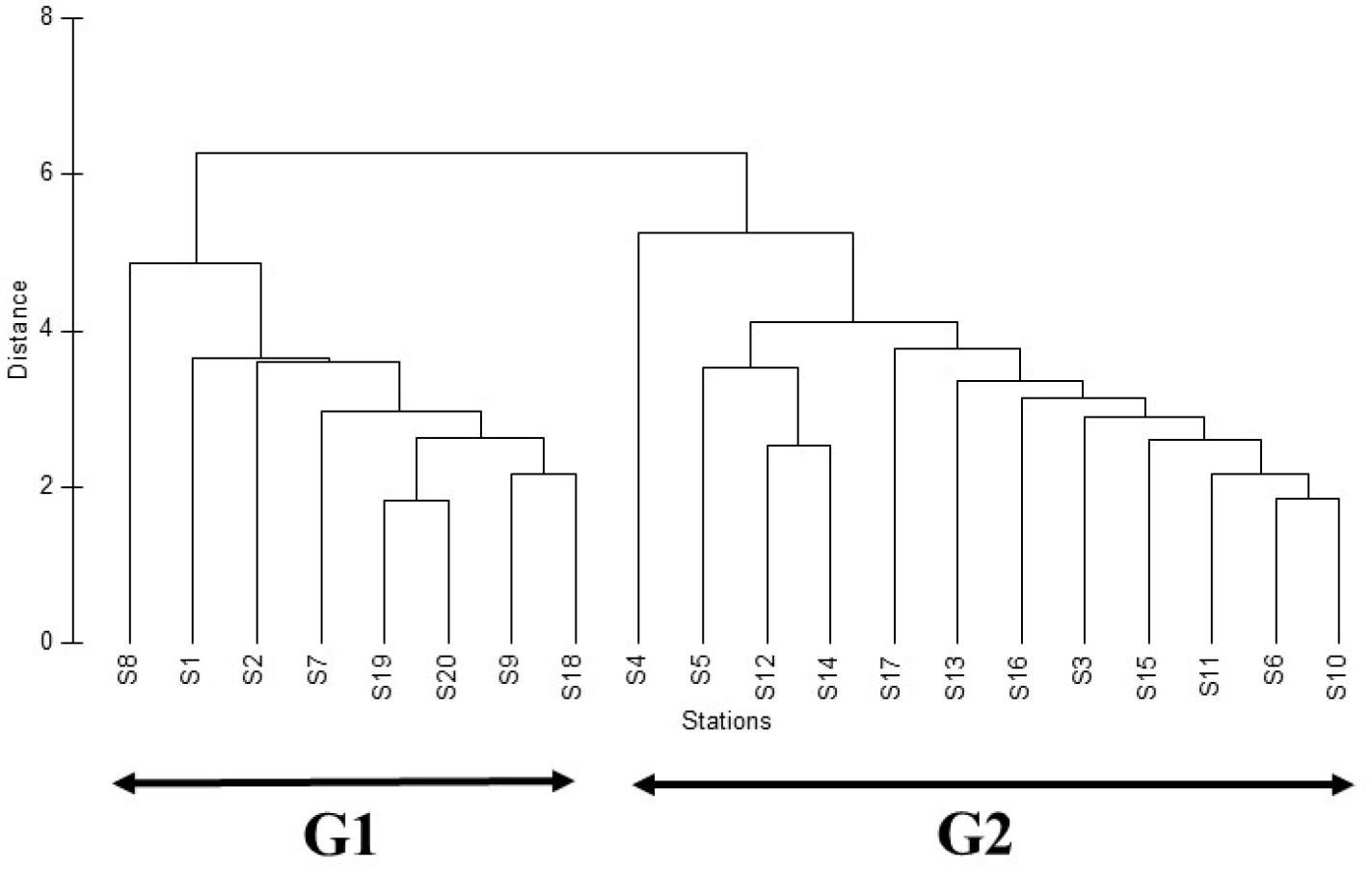
Cluster analysis based on *Gobius niger* abundances showing reciprocal relations between the 20 sampled stations in the Marchica lagoon according to Euclidean distance similarities.

DISTLM marginal tests assessed the importance of each variable separately. They indicated that depth, temperature and substratum had a significant effect (p<0.05), explaining respectively 49.6%, 22.5% and 21.3% of the spatial variations of *G. niger* in the Marchica lagoon (Table 1). Salinity and vegetation cover are close to the limit of significance and each explains about 13% of the total variance. The remaining variables, pH and suspended matter, did not show any significant impact and each accounted for less than 1% of the variability. The best model was obtained by a combination of depth, salinity, and suspended matter, which accounted for 58.89% of the variability in the data. In the dbRDA performed on the selected model, the first two axes accounted for 57.9% of the total variability (dbRDA1 = 56.6%; dbRDA2 = 1.3%) (Fig. 4). Depth was the main contributor to the first axis (loading dbRDA = 0.91), while the main contributors to the second factor were salinity and suspended matter (loading dbRDA = 0.57 and -0.64, respectively). Furthermore, the depth is negatively correlated with all parameters considered in the present study, with significance for temperature (Spearman’s test; r=-0.65, p= 0.001) and substrate (Spearman’s test; r=-0.52, p= 0.01).

**Table 1.** DISTLM marginal test results: Significance of the relationship between abiotic factors and monthly abundances of *Gobius niger*. p: p-value. Prop. (%): relative contribution of each environmental variable to variation in spatial structure.

| Variable | SS(trace) | Pseudo-F | p | Prop. (%) |
| --- | --- | --- | --- | --- |
| Temperature (Temp) | 63.356 | 5.2469 | 0.01* | 0.2257 |
| Depth | 139.49 | 17.781 | 0.001* | 0.49694 |
| Salinity | 38.621 | 2.8717 | 0.073 | 0.13759 |
| pH | 25.637 | 1.8092 | 0.138 | 9.13E-02 |
| Suspended Matter (SM) | 14 | 0.94485 | 0.348 | 4.99E-02 |
| VC (Vegetation Cover) | 36.035 | 2.651 | 0.072 | 0.12837 |
| MGS (Sediment Mean Grain Size) | 60.034 | 4.8969 | 0.012* | 0.21387 |

**Fig. 4:**
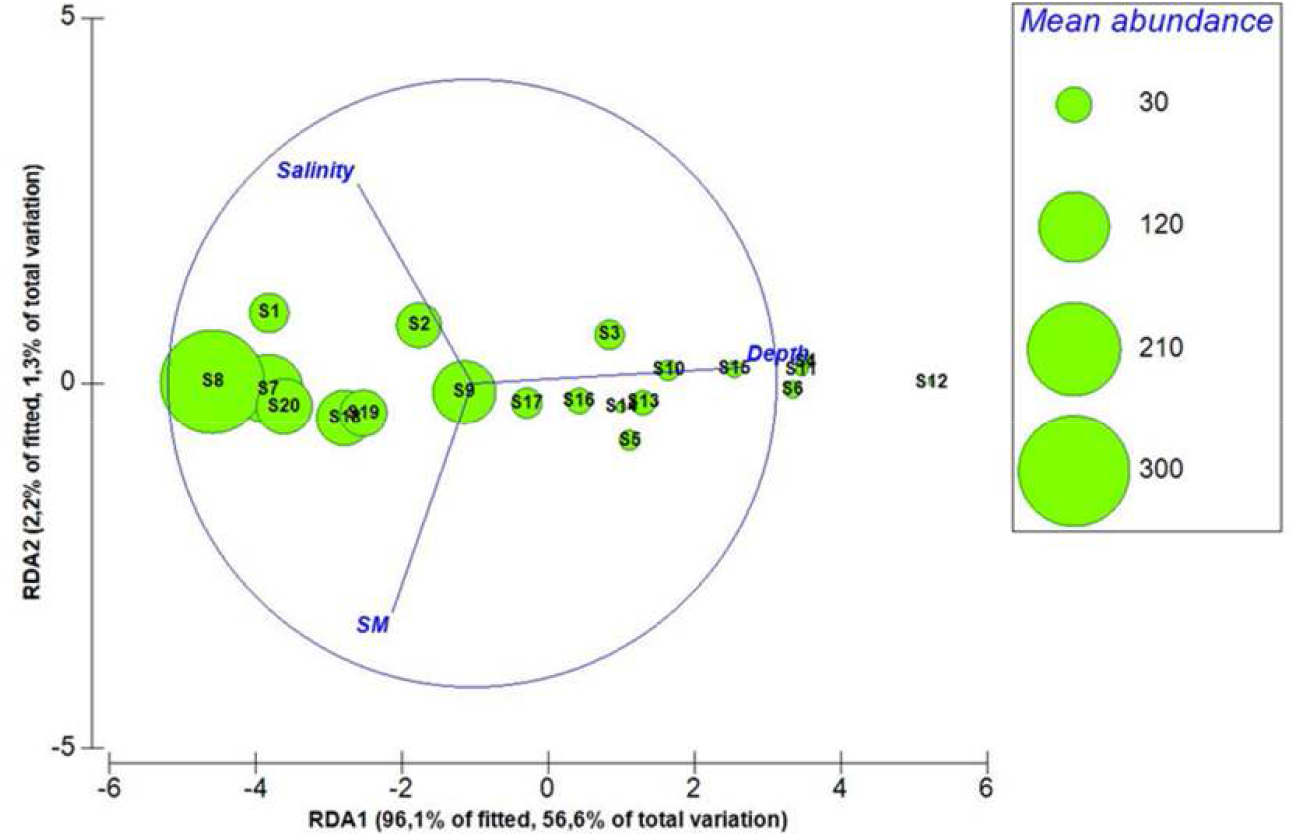
Two-dimensional redundancy analysis (RDA) ordination representing the model of spatial variation in spatial distribution of *Gobius niger* related to the predictor variables selected through the best linear models based on distance (DISTLM). SM: Suspended matter.

The black goby was present in the Marchica lagoon throughout the sampling year, both over space and time but with variable abundance (Fig. 5). Overall, the monthly abundance in all stations fluctuated between no individuals and a maximum of 643 individuals. The mean abundance (± SD) was 44.13 (± 88) individuals per station (Table S2). PERMANOVA results showed significant differences in total abundance between seasons and stations (Table 2). Interactions between the two factors were not significant (Pseudo F= 1.072, p (perm) > 0.05). A posteriori pairwise comparison revealed that the black goby was significantly less abundant in winter than in other seasons (Table 3). Regarding the second factor, most of the significant differences concern the combinations formed by the peripheral and central stations (Table S3).

**Fig. 5:**
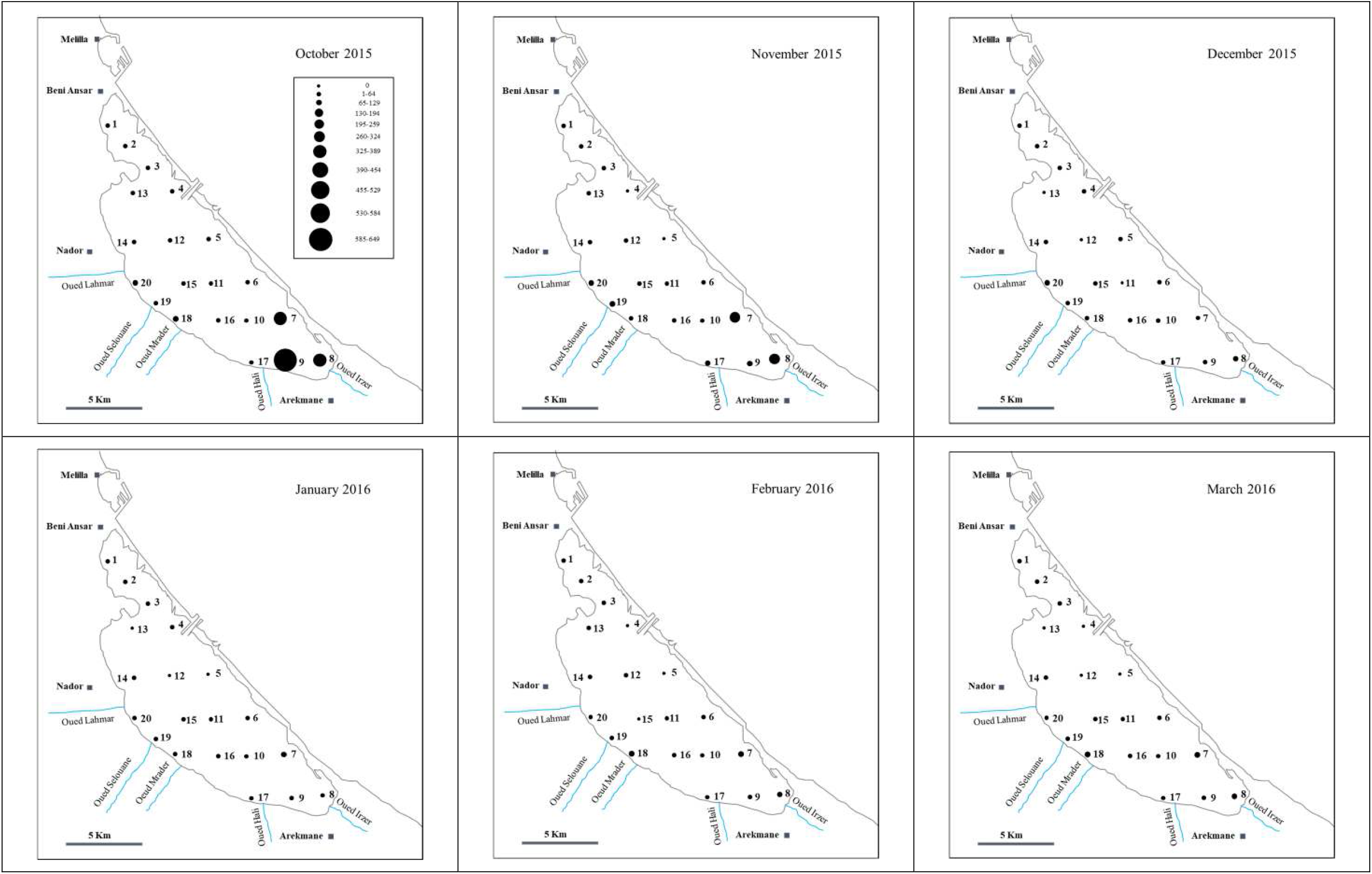
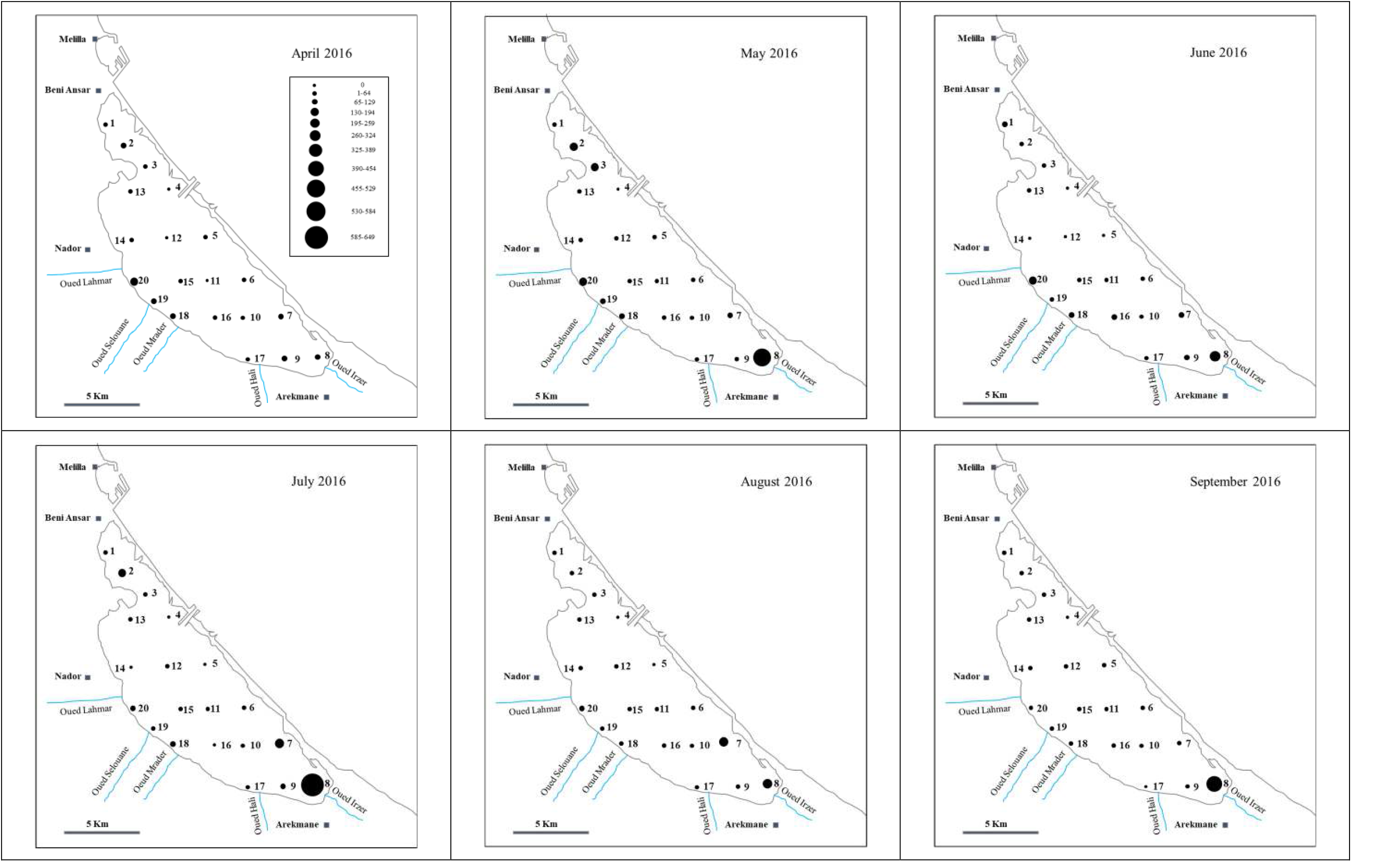
Spatial and temporal distribution of *Gobius niger* in the Marchica lagoon.

**Table 2.** Results of the multivariate permutational analysis (PERMANOVA) of Gobius niger considering season and station. Df: degrees of freedom; and p (perm): level of significance.

|  | Sum sq | Df | Mean sq | F value | Pperm |
| --- | --- | --- | --- | --- | --- |
| Season | 49908 | 3 | 16636 | 3.905 | 0.011* |
| Station | 860873 | 19 | 45309 | 10.636 | 0.001* |
| Season : Station | 260307 | 57 | 4567 | 1.072 | 0.302 |
| Residuals | 681627 | 160 | 4260 |  |  |

**Table 3.**
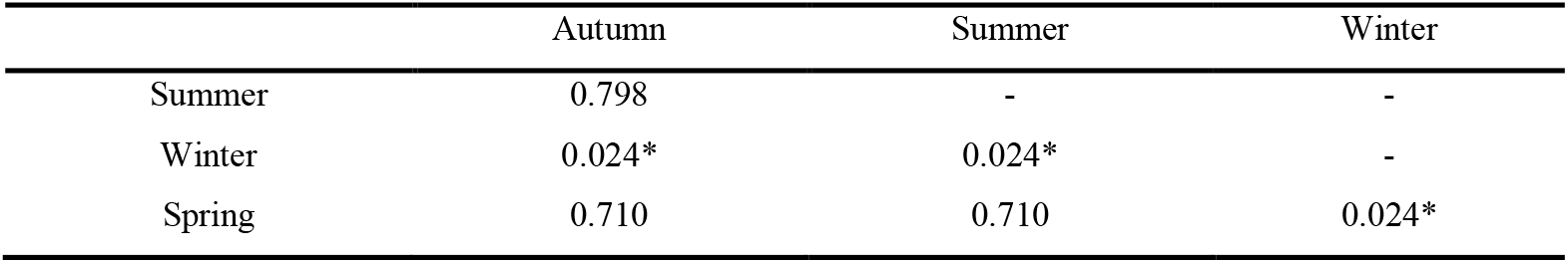
Pairwise comparisons of *Gobius niger* abundance in term of season.

### Characterisation and comparison of goby populations from the Marchica lagoon and adjacent Mediterranean Sea

#### Morphometric and meristic analysis

Data from meristic and morphometric characters of *Gobius niger* sampled at sea and in the Marchica lagoon are reported in Tables S4 to S7, respectively. Overall, specimens caught in the sea (109.1± 12 mm) differ significantly (p value < 0.05) from the ones caught the lagoon (85.4 ± 12.5 mm).

The allometric relationships (Log Y = a LogX = Logb) between the standard length (SL) and the total length (TL), the head length (HL) and the body height (BH), and on the other hand, between the head length (HL) and the eye diameter (ED) and the snout length (SnL) are summarized in Tables S8 and S9.

The first PCA (PC1 vs PC2) performed on the morphometric and meristic data allowed the identification of a distinct stratification based on the origin of the individuals. The first two axes are the most informative, representing respectively 78.67% and 9.75% of the total inertia, totalising 88.42% of the total variation. (Fig. 6); A plot of PC2 to PC3 shows no separation among these two groups (Fig. 6).

**Fig. 6:**
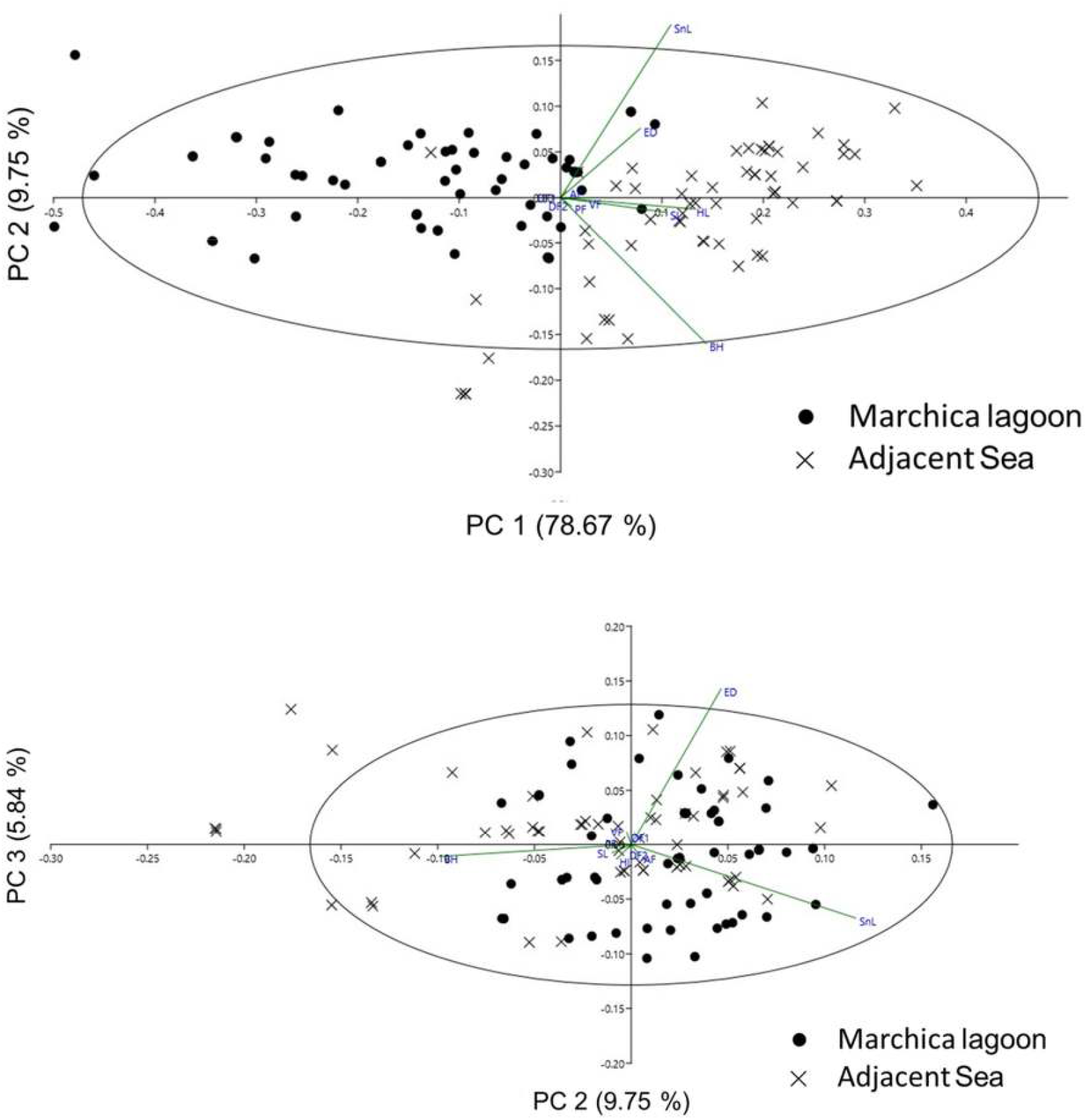
PCA of morphological variables of the *Gobius niger* (standard length, SL; body height, BH; head length, HL; snout length, SnL; eye diameter, ED; first dorsal fin, DF1; second dorsal fin, DF2; anal fin, AF; pectoral fin, PF and ventral fin, VF) with projection of phenotypic groups. PC1 vs. PC2 and PC2 vs. PC3. Percentage of variation explained by each PC axis is given within parenthesis.

Principal component 1 was strongly correlated to Standard length (Pearson Correlation 0.962) (Fig. 7). Principal components 2 and 3 were not correlated to SL (Pearson correlation -0.022 and -0.004 respectively) indicating that amongst-group differences on this axis are independent of specimen size and are interpretable directly as differences in shape.

**Fig. 7:**
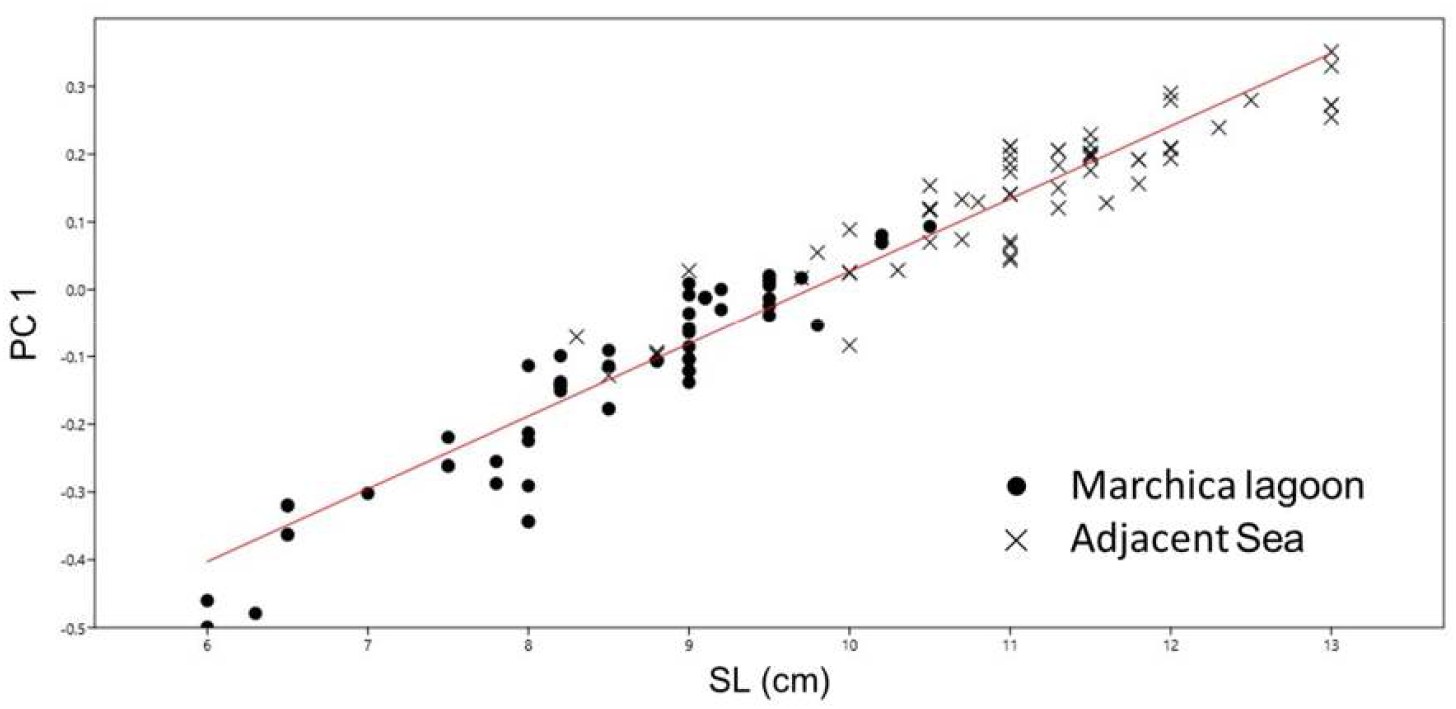
Linear regression of the principal component score axis (PC1) from morphometric measurements on the log standard length of *Gobius niger* with projection of phenotypic groups.

We can conclude that the morphometry is size depending, meristic is not, the second is expected, fin meristic can’t change with growth.

#### Genetic differentiation

From the 120 studied individuals of *G. niger* from Morocco, 88 sequences were obtained. The alignment of the portion of the 16S rRNA gene was 547 base pairs long and contained 26 polymorphic sites; a total of 31 haplotypes were found.

Diversity measures, calculated for the two groups of *Gobius niger* (Marchica lagoon and the adjacent sea), are reported in Table 4. The marine population shows 21 haplotypes while the population from the lagoon shows 17 haplotypes. The highest number of segregating sites (20) is recorded for the population of the adjacent sea while the population of the Marchica lagoon shows the lowest number of segregating sites (16). The nucleotide diversity (π) is 0.00413 for the group of the adjacent sea and 0.00462 for that of Marchica lagoon. The haplotype diversity for the adjacent sea is 0.85878 against 0.93741 in Marchica lagoon. Neutrality test values (Fu & Li’s F and Tajima’s D) are negative and significant.

**Table 4.**
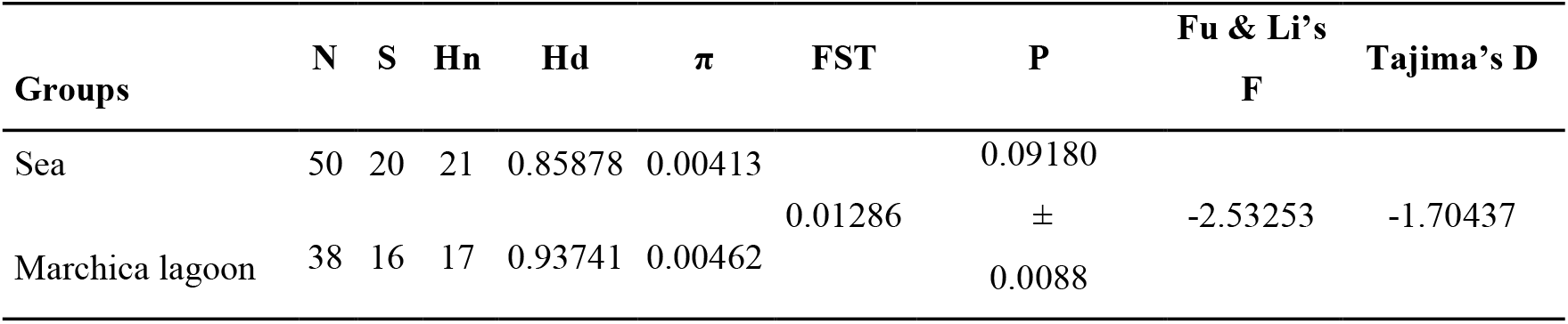
Results of the haplotype diversity analysis of the two groups of *Gobius niger*. N: number of sequences; S: number of segregating sites; Hn: number of haplotypes; Hd: haplotype diversity; π: nucleotide diversity; P: P-value. Significant values at α=0.05.

The pairwise F_ST_ comparisons shows low value; F_ST_ p-value is not significant. This implies that there is no considerable degree of genetic differentiation between the different groups of *G. niger* from Marchica lagoon, and the adjacent Mediterranean coast of Morocco.

The maximum parsimony network shows three ancestral haplotypes (numbers 22, 29, 41 in Fig. 8). There is no visible geographical segregation except for the sequences from Turkey and Greece whose haplotypes were not shared with the other individuals. The Western and Central Mediterranean and the Atlantic Ocean populations share the haplotypes, while the Eastern Mediterranean does not.

**Fig. 8:**
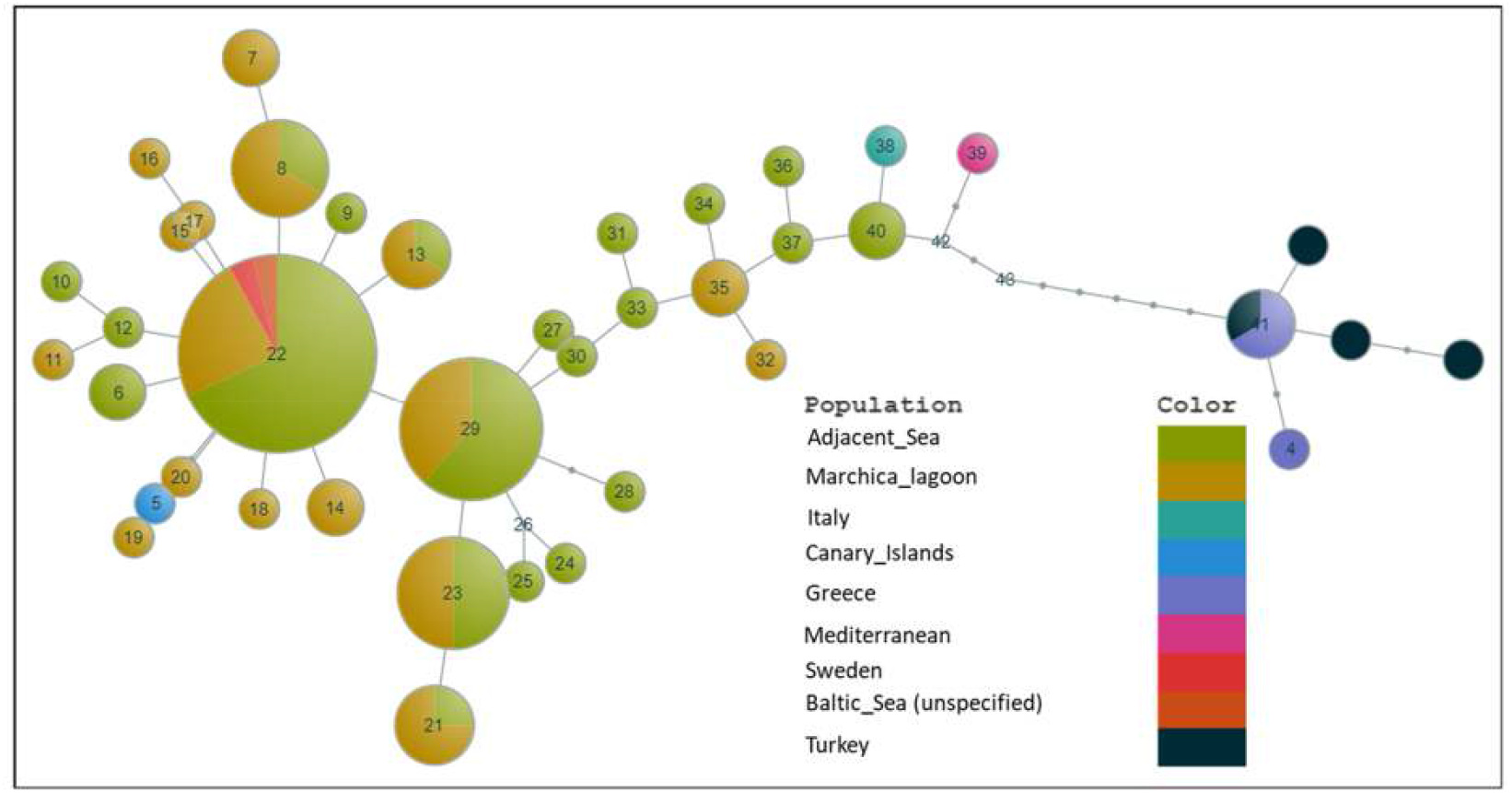
Haplotype network constructed from 16S rRNA sequences of *Gobius niger*. The size of a particular circle reflects the haplotype frequency. The numbers indicate the nodes.

The result of the Mantel test reports no correlation between the genetic and morphological distance matrices (r=-0.11; p=0.97), suggesting no isolation by distance and the existence of gene flow between the lagoon and the sea (Fig. 9).

**Fig. 9:**
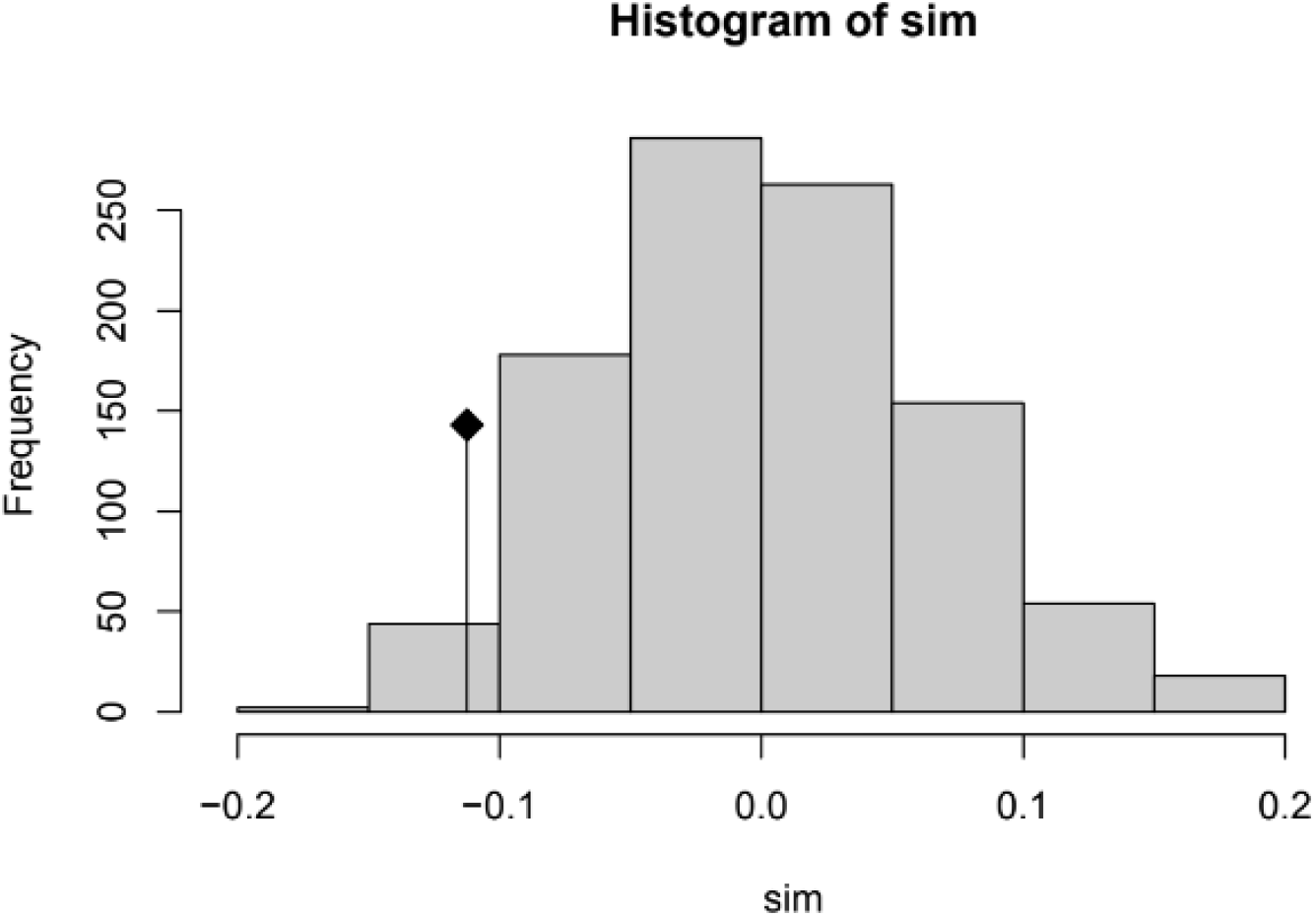
Histogram of the Mantel test assessing the relationship between genetic and morphologic distance for *Gobius niger*. Sim: simulations; Frequency: frequency values of the correlation between the genetic and morphologic distances; The dot represented the original value of the correlation between the distance matrices.

#### *Monogenean parasites from* G. niger

Of the 180 specimens of *G. niger* examined, no fish was parasitized by monogenean flatworms, either in the sea or in the Marchica lagoon.

## Discussion

### *Spatial and temporal distribution of* Gobius niger *in the Marchica lagoon and environmental drivers*

The spatial and temporal distribution of *G. niger* evidenced that the species is permanently present in the Marchica lagoon with higher abundances in the peripheral shallow bottoms of the lagoon on a variety of substrates (mud, muddy-sand, sandy-mud and fine sand) mostly covered by macroalgae and/or seagrass meadows (Najih *et al*., 2016, Najih *et al*., 2017). This is consistent with the ecology of the species in other Mediterranean coastal lagoons where it prefers shallow habitats (<1.5m depth) on sandy and sandy-muddy vegetated beds (Kara & Quignard, 2019). Moreover, its permanent and abundant presence in the Marchica lagoon supports resident species status (Franco *et al*.,2008a, 2008b, 2012, Kara & Quignard, 2019; Selfati *et al*., 2019) and very frequent fish (62.8%) in Mediterranean coastal lagoons (Kara & Quignard, 2019). *Gobius niger* plays a key eco-trophic role by providing a link between benthic invertebrates and large predators (Casabianca & Kiener, 1969; Miller, 1979; Raffaelli *et al*., 1989). Therefore, the shallow beds of the lagoon, where the species is abundant, are key habitats in the Marchica lagoon and need to be considered in all management plans aiming at the conservation of biodiversity and ecological processes. Moreover, being a benthic and resident fish, *G. niger* is a suitable candidate to monitor the ecological status of the Marchica lagoon, especially in its most disturbed peripheral areas (Ben Hassine *et al*., 1999). Indeed, *G. niger* has been used in many pollution monitoring studies (Fossi *et al*., 1989; Katalay & Parkak, 2002; Migliarini *et al*., 2005; Maradonna *et al*., 2004; Barucca *et al*., 2006; Barucca *et al*., 2006; Maradonna & Carnevali, 2007; Ramsak *et al*., 2007).

Overall, depth seems to be the most important predictor variable, explaining the distribution of *G. niger* in the lagoon. This is in accordance with the distribution patterns of fish assemblages in the Marchica lagoon where a spatial gradient in the bentho-demersal component of the fish assemblage structure was observed, with more marine species occurring near the sea inlet and more resident species in the inner margins of the lagoon (Selfati *et al*., 2019). Moreover, depth is a key factor for understanding the structure of fish assemblages in the Marchica lagoon, but has to be seen as a variable that acts in concert with other factors such as vegetation cover, pH, temperature, dissolved oxygen, salinity and suspended matter (Selfati *et al*. 2019). Furthermore, the depth is negatively correlated with all parameters considered in the present study, with significance for temperature (Spearman’s test; r=-0.65, p= 0.001) and type of substrate (Spearman’s test; r=-0.52, p= 0.01). This assumes that the shallower areas of the lagoon present differential conditions in terms of temperature, luminosity and other conditions to support primary productivity. This is in line with the ecological requirements of the black goby, particularly in terms of reproduction, feeding and shelter from predation. Black goby requires coarse substrates to have shelters for nest construction (Vaas *et al*. 1975; Wiederholm, 1987; Mistri *et al*., 2000; Leatemia *et al*., 2017). The vegetated habitats are important for the abundance of marine macroinvertebrates which are the main prey resource for the black goby (Hajji *et al*., 2013; Matern *et al*., 2021). Vegetation cover also limits predation by increasing the complexity of the habitats (Heck and Orth, 1980; Kulczycki *et al*., 1981). The significant decrease of *G. niger* during winter could be related to low temperatures during this season leading to migration of the species to deeper marine waters (Mozzo, 1968).

### Characterisation and comparison of goby populations from the Marchica lagoon and adjacent Mediterranean Sea

#### Morphometric patterns

In the Moroccan Mediterranean, the black goby has been previously reported both in the lagoon and in the open sea (Aloncle, 1951; Selfati *et al*., 2019). The work of Böhlke & Robins (1968), Hoese (1971, 1983), Bath (1973), Akihito (1986), Gill *et al*., (1992), and more recently by Bouchereau *et al*. (2000) and Kovačić & Golani (2007) has shown that the standard length (SL) and head length (HL) are very useful in the classification of genera and species of Gobiidae. The difference in size between individuals from the lagoon and their marine counterparts is probably due to the depth difference of the two sites. Indeed, the majority of fishes show a size positively correlated with depth (Harvey & Stewart, 1991). Fish are exposed to the risk of predation; to reduce it, they choose deeper habitats, and therefore will have a longer life span which should be reflected to some extent by a larger body size (Harvey & Stewart, 1991).

#### Genetic structure: lagoon and open sea

The genetic structure of marine fish populations reflects the historical and contemporary interaction between a complex set of ecological, demographic, behavioral, genetic, oceanographic, climatic, and tectonic processes. The combined effect of these mechanisms, acting on a range of spatial and temporal scales, determines the rates and patterns of dispersal of gametes, zygotes, larvae and adults (Giovannotti *et al*., 2009).

Our study revealed the presence of common haplotypes shared by two groups from different habitats. According to Bortone *et al*. (2005), *G. niger* is considered as a resident species in the Marchica lagoon (Selfati, 2020). However, the results of our study suggest recurrent migrations between the groups of the sea and lagoon. Indeed, the haplotype network reconstruction revealed no apparent population subdivision and no geographical segregation between lagoon and sea. F_ST_ values are used to conclude on the level of gene flow (Chanthran *et al*., 2020). According to Wright (1965), populations with F_ST_ of 0–0.05 show a small differentiation, 0.05–0.15 is considered as moderate differentiation, 0.15–0.25 as important differentiation and values greater than 0.25 as very important differentiation. The occurrence of common haplotypes between the groups and small to moderate differentiation based on F_ST_ results suggests unrestricted gene flow with no relation to the habitat.

#### *Parasites of* G. niger

Monogenean parasites are one of the largest groups of Platyhelminthes characterised by high species diversity and high host specificity (Gusev, 1995; Kearn, 1994; Poulin, 1998). The most abundant genus of monogeneans in gobies as well as in many other fishes is *Gyrodactylus* von Nordmann, 1832. According to Huyse and Volckaert (2005) species of *Gyrodactylus* parasitize on 19 orders of fresh and marine bony fishes. To date, three *Gyrodactylus* spp. parasitizing *G. niger* have been recorded: *Gyrodactylus flesi* Malmberg, 1957, *G. proterorhini* Ergens, 1967 and *G. niger* Huyse *et al*. 2003 from North Sea (Huyse *et al*., 2003; Harris *et al*., 2004).

The absence of monogenean parasites on the gills and the skin of *G. niger* from the Marchica lagoon and sea could be explained by environmental conditions. Another explanation of this absence is the immune system of fishes. Indeed, Zander (1993) and Zander *et al*. (1999) showed that in the Baltic Sea, *Podocotyle atomon* (Rudolphi, 1802) is present in high abundances; however, in *G. niger* this species was often absent, which was explained by its immune system efficiency.

## Conclusion

*Gobius niger* was revealed to be present year-round in the Marchica lagoon with higher abundances recorded in the shallow bottoms of the lagoon inner margins on a variety of mostly vegetated substrates and with general decrease in abundances during winter. Depth is a key factor for understanding the spatial patterns of *G. niger* in Marchica lagoon, but has to be seen as a variable that acts in concert with other factors such as vegetation cover, type of substrate and temperature among others. Thus, the shallow beds of the lagoon, where the species is abundant, are key habitats in the Marchica lagoon and need to be considered in all management plans aiming at the conservation of biodiversity and ecological processes. Comparison of black goby populations from the Marchica lagoon with their counterparts from the adjacent Mediterranean coast of Morocco revealed that specimens caught in the sea are a significantly bigger compared to the ones from the lagoon. Moreover, the absence of population structuring and common haplotypes between the two populations indicates no apparent restriction in the gene flow between the two populations.

## Acknowledgements

This research was supported by the Special Research Fund of Hasselt University (BOF23BL07 to A.L.; BOF21PD01 to N.K.; BOF20TT06 to M.P.M.V.; BOF21INCENT09) and by research grant 1513419N of the Research Foundation – Flanders (FWO-Vlaanderen). The sampling survey was undertaken in the framework of an international cooperation between Le Conservatoire du Littoral, Agence de l’Eau Rhône-Méditerranée-Corse, the Ecocean Society, University Mohammed V in Rabat, Institut National de Recherche Halieutique, Observatoire de la Marchica, and Fondation Mohammed VI pour l’environnement. M. Selfati thanks the “Agence de l’Eau Rhône Méditerranée Corse” and the Ecocean Society for their financial support. Authors are very grateful to fishers for providing *Gobius niger* samples from both the Marchica lagoon and the adjacent Mediterranean coast of Morocco. Dr. Lukas Rüber (Natural History Museum of Bern, Switzerland) is thanked for curatorial services.

## Supplementary material

The following supplementary material is available for this article:

**Figure S1.**
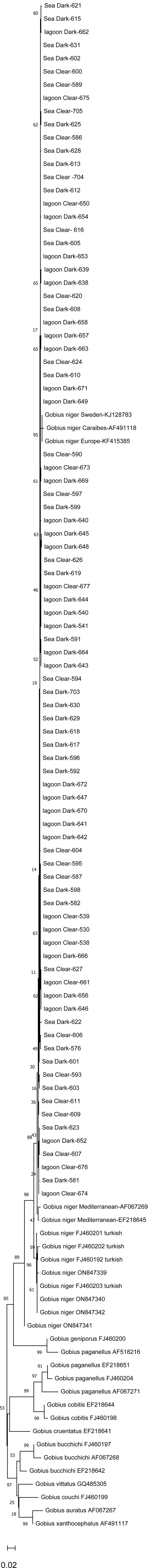
A neighbour-joining tree including all the species of *Gobius* that are represented on GenBank by a targeted fragment of the 16S rRNA gene. The tree was created in MEGA X using 1000 bootsrap replicas and the kimura 2-P substitution model. the scale bar represents the number of expected substitutions per site.

**Table S1.** Summary of species included in the phylogenetic tree, with accession numbers and locations of each species.

| Species | Accession number | Locations | References |
| --- | --- | --- | --- |
| *Gobius niger* | KJ128783 | Sweden | Ericson, 2020 |
|  | KF415385 | Baltic Sea (unspecified) | Agoretta, 2013 |
|  | EF218645 | Italy, Ancona | Giovannotti, 2007 |
|  | AF067269 | Mediterranean | Penzo, 1998 |
|  | FJ460203 | Turkey | Turan, 2016 |
|  | FJ460202 | Turkey | Turan, 2016 |
|  | FJ460201 | Turkey | Turan, 2016 |
|  | FJ460192 | Turkey | Turan, 2016 |
|  | AF491118 | Canary Islands | Ruber, 2003 |
|  | ON847339 | Greece |  |
|  | ON847340 | Greece |  |
|  | ON847341 | Greece |  |
|  | ON847342 | Greece |  |
| *Gobius paganellus* | AF518216 | Unpublished |  |
|  | EF218651 | Italy, Ancona | Giovannotti, 2007 |
|  | FJ460204 | Turkey | Turan, 2016 |
|  | AF067271 | Mediterranean | Penzo, 1998 |
| *Gobius bucchichi* | EF218642 | Italy, Napoli | Giovannotti, 2007 |
|  | FJ460197 | Turkey | Turan, 2016 |
|  | AF067268 | Mediterranean | Penzo, 1998 |
| *Gobius cobitus* | EF218644 | Italy, Ancona | Giovannotti, 2007 |
|  | FJ460198 | Turkey | Turan, 2016 |
| *Gobius cruentatus* | EF218641 | Italy, Napoli | Giovannotti, 2007 |
| *Gobius auratus* | AF067267 | Mediterranean | Penzo, 1998 |
| *Gobius xanthocephalus* | AF491117 | Spain, Gran Canaria, Puerto Rico | Ruber, 2003 |
| *Gobius vitatus* | GQ485305 | Turkey | Turan, 2016 |
| *Gobius couchi* | FJ460199 | Turkey | Turan, 2016 |
| *Gobius geniporus* | FJ460200 | Turkey | Turan, 2016 |

**Table S2.** Mean abundance of the *Gobius niger* species at different stations.

|  | S1 | S2 | S3 | S4 | S5 | S6 | S7 | S8 | S9 | S10 | S11 | S12 | S13 | S14 | S15 | S16 | S17 | S18 | S19 | S20 |
| --- | --- | --- | --- | --- | --- | --- | --- | --- | --- | --- | --- | --- | --- | --- | --- | --- | --- | --- | --- | --- |
| **October** | 41 | 3 | 1 | 17 | 1 | 2 | 345 | 470 | 610 | 20 | 4 | 1 | 9 | 5 | 3 | 30 | 18 | 92 | 32 | 70 |
| **Novomber** | 37 | 21 | 1 | 0 | 0 | 3 | 150 | 130 | 28 | 29 | 5 | 1 | 2 | 1 | 2 | 16 | 60 | 58 | 70 | 116 |
| **December** | 22 | 6 | 10 | 7 | 2 | 1 | 63 | 88 | 20 | 1 | 0 | 0 | 0 | 1 | 8 | 1 | 4 | 9 | 40 | 66 |
| **January** | 11 | 14 | 1 | 49 | 0 | 6 | 90 | 52 | 12 | 7 | 1 | 0 | 0 | 1 | 1 | 3 | 14 | 13 | 51 | 30 |
| **February** | 2 | 16 | 9 | 0 | 0 | 24 | 67 | 103 | 60 | 3 | 4 | 1 | 18 | 7 | 0 | 5 | 17 | 80 | 57 | 40 |
| **March** | 10 | 26 | 5 | 0 | 0 | 5 | 113 | 76 | 56 | 3 | 1 | 0 | 0 | 3 | 5 | 7 | 20 | 102 | 60 | 30 |
| **April** | 6 | 93 | 8 | 0 | 1 | 1 | 74 | 116 | 78 | 3 | 0 | 0 | 5 | 3 | 1 | 15 | 26 | 110 | 108 | 144 |
| **May** | 12 | 176 | 143 | 0 | 62 | 5 | 66 | 494 | 49 | 6 | 9 | 1 | 22 | 3 | 15 | 29 | 8 | 87 | 103 | 113 |
| **June** | 130 | 6 | 3 | 0 | 0 | 1 | 9 | 300 | 69 | 4 | 5 | 0 | 50 | 0 | 10 | 68 | 15 | 100 | 44 | 90 |
| **July** | 36 | 174 | 5 | 0 | 0 | 3 | 196 | 643 | 121 | 9 | 2 | 0 | 14 | 0 | 3 | 0 | 36 | 190 | 11 | 90 |
| **August** | 43 | 6 | 56 | 0 | 0 | 26 | 190 | 230 | 24 | 19 | 8 | 2 | 4 | 5 | 5 | 2 | 45 | 19 | 27 | 80 |
| **September** | 94 | 37 | 6 | 0 | 53 | 7 | 42 | 430 | 25 | 19 | 3 | 3 | 48 | 1 | 27 | 9 | 0 | 41 | 36 | 43 |
| **Mean abundance** | 37 | 48 | 21 | 6 | 10 | 7 | 117 | 261 | 96 | 10 | 4 | 1 | 14 | 3 | 7 | 15 | 22 | 75 | 53 | 76 |

**Table S3.** Pairwise comparisons of *Gobius niger* abundance in term of factor station.

|  | S1 | S10 | S11 | S12 | S13 | S14 | S15 | S16 | S17 | S18 | S19 | S2 | S20 | S3 | S4 | S5 | S6 | S7 | S8 |
| --- | --- | --- | --- | --- | --- | --- | --- | --- | --- | --- | --- | --- | --- | --- | --- | --- | --- | --- | --- |
| S10 | 0.0292 | - | - | - | - | - | - | - | - | - | - | - | - | - | - | - | - | - | - |
| S11 | 0.0051 | 0.0577 | - | - | - | - | - | - | - | - | - | - | - | - | - | - | - | - | - |
| S12 | 0.0051 | 0.0051 | 0.0086 | - | - | - | - | - | - | - | - | - | - | - | - | - | - | - | - |
| S13 | 0.1237 | 0.5689 | 0.0577 | 0.0086 | - | - | - | - | - | - | - | - | - | - | - | - | - | - | - |
| S14 | 0.0051 | 0.0200 | 0.4947 | 0.0577 | 0.0292 | - | - | - | - | - | - | - | - | - | - | - | - | - | - |
| S15 | 0.0086 | 0.3875 | 0.3024 | 0.0051 | 0.2447 | 0.0848 | - | - | - | - | - | - | - | - | - | - | - | - | - |
| S16 | 0.1366 | 0.5689 | 0.0233 | 0.0051 | 0.9551 | 0.0200 | 0.2379 | - | - | - | - | - | - | - | - | - | - | - | - |
| S17 | 0.3055 | 0.0823 | 0.0051 | 0.0051 | 0.3923 | 0.0051 | 0.0292 | 0.4906 | - | - | - | - | - | - | - | - | - | - | - |
| S18 | 0.0966 | 0.0051 | 0.0051 | 0.0051 | 0.0051 | 0.0051 | 0.0051 | 0.0051 | 0.0051 | - | - | - | - | - | - | - | - | - | - |
| S19 | 0.3162 | 0.0051 | 0.0051 | 0.0051 | 0.0051 | 0.0051 | 0.0051 | 0.0086 | 0.0086 | 0.2635 | - | - | - | - | - | - | - | - | - |
| S2 | 0.6915 | 0.0766 | 0.0051 | 0.0051 | 0.1257 | 0.0051 | 0.4585 | 0.1608 | 0.2863 | 0.3445 | 0.8170 | - | - | - | - | - | - | - | - |
| S20 | 0.3912 | 0.0051 | 0.0051 | 0.0051 | 0.0051 | 0.0051 | 0.0051 | 0.0051 | 0.0051 | 0.9551 | 0.1561 | 0.2726 | - | - | - | - | - | - | - |
| S3 | 0.3875 | 0.6116 | 0.1134 | 0.0051 | 0.8398 | 0.0523 | 0.4585 | 0.8088 | 0.9551 | 0.0233 | 0.0523 | 0.2950 | 0.0125 | - | - | - | - | - | - |
| S4 | 0.0266 | 0.5216 | 0.8088 | 0.2515 | 0.2830 | 0.6415 | 0.9551 | 0.2982 | 0.0650 | 0.0051 | 0.0051 | 0.0358 | 0.0051 | 0.2988 | - | - | - | - | - |
| S5 | 0.0650 | 0.9551 | 0.5642 | 0.4677 | 0.6556 | 0.5642 | 0.7249 | 0.6116 | 0.2515 | 0.0086 | 0.0051 | 0.1112 | 0.0051 | 0.5512 | 0.7492 | 0.7492 | - | - | - |
| S6 | 0.0086 | 0.5154 | 0.3599 | 0.0086 | 0.3019 | 0.1216 | 0.9560 | 0.2796 | 0.326 | 0.0051 | 0.0051 | 0.0292 | 0.0051 | 0.4906 | 0.9560 | 0.7405 | - | - | - |
| S7 | 0.0233 | 0.0051 | 0.0051 | 0.0051 | 0.0051 | 0.0051 | 0.0051 | 0.0051 | 0.0051 | 0.2467 | 0.0200 | 0.0741 | 0.2476 | 0.0051 | 0.0051 | 0.0086 | 0.0051 | - | - |
| S8 | 0.0051 | 0.0051 | 0.0051 | 0.0051 | 0.0051 | 0.0051 | 0.0051 | 0.0051 | 0.0051 | 0.0051 | 0.0051 | 0.0086 | 0.0125 | 0.0051 | 0.0051 | 0.0051 | 0.0051 | 0.0605 | - |
| S9 | 0.2761 | 0.0086 | 0.0051 | 0.0051 | 0.0125 | 0.0051 | 0.0051 | 0.0086 | 0.0266 | 0.9479 | 0.7272 | 0.5642 | 0.9560 | 0.0650 | 0.0051 | 0.0086 | 0.0051 | 0.8008 | 0.0650 |

**Table S4.**
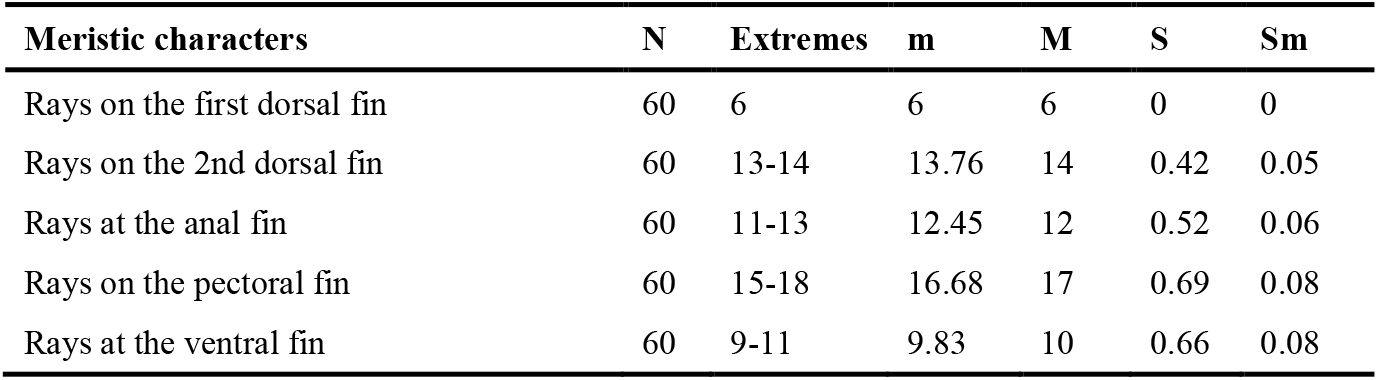
Meristic characters of *Gobius niger* from the Mediterranean Sea. N: number; m: mean; M: mode; S: standard deviation; Sm: standard error of the mean.

**Table S5.**
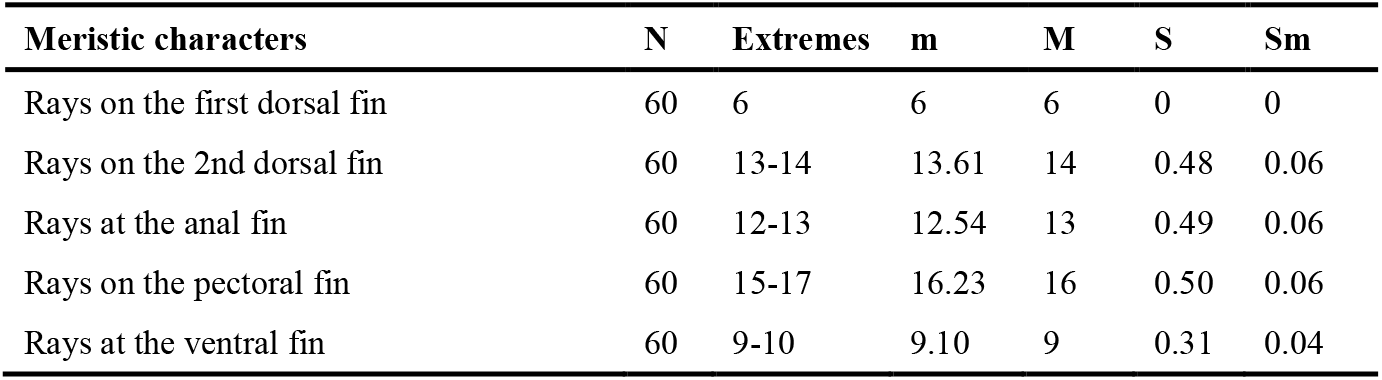
Meristic characters of *Gobius niger* from Marchica lagoon. N: number; m: mean; M: mode; S: standard deviation; Sm: standard error of the mean.

**Table S6.** Morphometric characters of *Gobius niger* from the Mediterranean Sea. N: number; m: mean; M: mode; S: standard deviation; Sm: standard error of the mean.

| Morphometric characters | N | Extremes | m | M | S | Sm |
| --- | --- | --- | --- | --- | --- | --- |
| Total length | 60 | 9-16 | 13.25 | 14 | 1.51 | 0.20 |
| Standard length | 60 | 8-13.5 | 10.91 | 11 | 1.20 | 0.16 |
| Head length | 60 | 2.3-4 | 3.08 | 3 | 0.41 | 0.05 |
| Snout length | 60 | 0.6-1.5 | 1.05 | 1 | 0.20 | 0.03 |
| Body height | 60 | 1.5-3.1 | 2.50 | 2.5 | 0.28 | 0.04 |
| Eye diameter | 60 | 0.3-0.5 | 0.41 | 0.4 | 0.05 | 0.01 |

**Table S7.** Morphometric characters of *Gobius niger* from the Marchica lagoon. N: number; m: mean; M: mode; S: standard deviation; Sm: standard error of the mean.

| Morphometric characters | N | Extremes | m | M | S | Sm |
| --- | --- | --- | --- | --- | --- | --- |
| Total length | 60 | 6.3-13 | 10.45 | 11.5 | 1.59 | 0.22 |
| Standard length | 60 | 5.4-10.5 | 8.54 | 9 | 1.25 | 0.17 |
| Head length | 60 | 1.5-3 | 2.30 | 2.5 | 0.40 | 0.06 |
| Snout length | 60 | 0.5-1.2 | 0.88 | 1 | 0.17 | 0.02 |
| Body height | 60 | 0.8-2.4 | 1.72 | 2 | 0.36 | 0.05 |
| Eye diameter | 60 | 0.3-0.4 | 0.32 | 0.4 | 0.04 | 0.01 |

Table S8. Allometric relationships between various numerical characters measured in *Gobius niger* from the Mediterranean Sea. SL: standard length; TL: total length; HL: head length; BH: body height; ED: eye diameter, N: number; r: correlation coefficient.

Table S9. Allometric relationships between various numerical characters measured in *Gobius niger* from the Marchica lagoon. SL: standard length; TL: total length; HL: head length; BH: body height; ED: eye diameter, N: number; r: correlation coefficient.

